# A Comparison of Macaque Hair Hormone Concentration Following Enhanced Cognitive Experiences or Standard Nonhuman Primate Environmental Enrichment

**DOI:** 10.1101/2021.12.01.470773

**Authors:** Brooke J. Meidam, Emilia K. Meredith, Amita Kapoor, Allyson J. Bennett, Peter J. Pierre

## Abstract

Experience with enriched environments positively impacts the health and wellbeing of nonhuman animals ranging from rodents to primates. Little is known, however, about the specific effects of enhanced cognitive enrichment (ECE) on nonhuman primates. The study reported here used archival samples to provide preliminary analysis of ECE on hormones associated with stress and wellbeing, as well as evaluation of persistent effects of infant social rearing. Hair samples from 24 adult male rhesus macaques were analyzed via LC-MS/MS technique for the main stress response hormones: cortisol, cortisone, and dehydroepiandrosterone. The ECE and care-as-usual (CAU) groups did not significantly differ in any of the mean analyte values. Cortisol and dehydroepiandrosterone were strongly and positively correlated in the ECE group. By contrast, for the CAU group the two analytes were weakly correlated. Within the ECE group, cortisol and dehydroepiandrosterone were significantly and positively correlated in the mother-reared group, but not in the nursery-reared group. Overall, these results provide preliminary evidence that experience with ECE could help to maintain a balanced cortisol:dehydroepiandrosterone ratio, possibly indicative of a healthy stress response. Further examination of this ratio in hair is needed to support this hypothesis. These observations may also suggest that nursery rearing could have persistent effects, including dysregulation of the hypothalamic-pituitary-adrenal axis, apparent in the unbalanced cortisol:dehydroepiandrosterone ratio. Together these findings are consistent with the growing literature that supports the use of ECE to promote nonhuman primate wellbeing and healthy development.

## 1. INTRODUCTION

Decades of research with laboratory-housed animals has demonstrated that enriched environments (EE) result in biological, neurological, and psychological changes (Renner & Rosenzweig, 1987). Currently, environmental enrichment is provided to a wide range of research animals, with the goal of promoting species typical behavior and psychological wellbeing. The effects of EE in rodents have been studied since the 1940’s (Tryon, 1940) and since this time, a large body of evidence has been generated which demonstrates the various beneficial effects of EE on health and wellbeing in rodents (for review see Renner & Rosenzweig, 1987; Olsson & Dahlborn, 2002). More recent studies have demonstrated that the effects physical enrichment in rats can differ based on social environment. For example, Zaias et al. (2008) found that rats in an impoverished environment ate more food, exhibited more locomotor activity and had greater body weights than their enriched counterparts. The researchers also found an interaction between the physical and social enrichment variables, such that the housing group size only had an effect in the physically enriched conditions. As the literature has grown, care practices have been adapted to reflect new knowledge, allowing extensive studies of physiological, neural, behavioral, immunological and other outcomes with direct comparisons of animals with EE and controls (Benefiel et al., 2005; Bennett et al., 1964; Diamond et al., 1964; Fischer, 2016; Frasca et al., 2013; Hebb, 1949; for review see Renner & Rosenzweig, 1987; Simpson & Kelly, 2011). Despite a large body of literature on EE in NHP, however, relatively less is known due to fewer controlled studies with neurobiological and physiological measurements aimed at evaluating the breadth of biological and psychological changes that may occur with differential exposure to EE in nonhuman primates (NHP). Provision of environmental enrichment for NHP has been a legal requirement since 1985 (Animal Welfare Act, 1985; Title 9C.F.R., Chapter 1, Subchapter A—Animal Welfare, Parts 3, Section 3.81). Subsequently, much research has been published regarding what constitutes beneficial enrichment for NHP. Considerable research has focused on the effects of various physical enrichment practices on common behavioral measurements of wellbeing (i.e. reactivity, increase in species typical behavior, decreases in abnormal behavior). According to a 2014 survey, common physical enrichment practices typically consist of nonresponse-contingent enrichment devices that promote manipulation or extraction of food, such as foraging devices or puzzle feeders (Baker, 2016). While these care-as-usual (CAU) practices typically elicit short term positive behavioral changes, many of these practices lack evidence for eliciting sustained engagement or providing a cognitive challenge for NHP (see Bennett et al., 2018 for additional review; Bennett et al., 2014; Washburn, 2015). Decades of research demonstrating intrinsic curiosity and exploratory drives in NHP suggest that providing opportunities for learning through cognitive challenges is of high importance for NHP wellbeing. There is little evidence that current enrichment practices address these drives. Provision of enhanced cognitive enrichment (ECE) through videogames or automated testing systems is a way to offer response-contingent, cognitively challenging enrichment for captive NHP that may satisfy their intrinsic curiosity and exploratory drives, thereby increasing NHP wellbeing.

Previous studies by our lab and others have provided ample evidence that experience with ECE through provision of videogames and other automated testing systems is beneficial to NHP health and wellbeing. It has even been argued previously that computer-based testing is an effective form of environmental enrichment which promotes psychological wellbeing in NHP, as access to computerized testing resulted in a higher wellbeing index regardless of housing status (Washburn, 2015). Videogames and automated testing systems elicit sustained use by NHP, reduce stereotypic behaviors and time spent resting, and increase locomotion and social behavior (Bennett et al., 2016; Fagot et al., 2014). Further, several studies provide evidence that nonhuman primates choose video games over free video and that use of the devices is insensitive to housing status, system parameters, and reward received (i.e. video, juice, or pellet) (Andrews & Rosenblum, 1993; Calapai et al., 2017; Washburn et al., 1994; Washburn & Hopkins, 1994; Washburn & Rumbaugh, 1992b). These studies primarily operationalize benefit to the animals in terms of amount of manipulation of the device or evaluate wellbeing using behavioral measures.

To our knowledge, only one study has evaluated the effect of experience with ECE on physiological measures of wellbeing. A study by Fagot and colleagues (2014), in baboons, found that salivary cortisol levels were lower after using an automated testing system. Glucocorticoids are commonly used measurements of wellbeing in primate studies that are indicative of stress level because they reflect the functioning of the HPA axis, which is activated in response to stressors (Nicolaides et al., 2015). Cortisol is the most abundant glucocorticoid in primates and can be measured in blood, saliva, or hair. Hair cortisol has become a common, noninvasive measurement of HPA axis functioning. Its utility lies in its ability to assess a more chronic state as compared to acute measurement from a blood sample. Hormones in hair are thought to reflect endocrine activity over a period of one to three months (Novak et al., 2017). This timeline makes hair an ideal measurement of baseline HPA axis activity during periods when stress is not experienced by the animal and in turn, a baseline measurement of wellbeing. In fact, evidence from primates demonstrate that hair cortisol concentrations increase after periods of stress (Zijlmans et al., 2021) and decrease after the stressor is removed (Davenport et al., 2006). In humans, both chronically high and low cortisol levels are associated with poor health and affective disorders such as anxiety, depression, and post-traumatic stress disorder (Fries et al., 2005; see Sharpley et al., 2011 for additional review; Sriram et al., 2012). Together the evidence suggest that hair cortisol values may be reflective of an animal’s state of wellbeing, providing a useful physiological measure of the effectiveness of care and enrichment practices.

In addition to cortisol, research on hormones in human hair shows that follicles synthesize 11-beta hydroxysteroid dehydrogenase (11β-HSD), an enzyme that converts cortisol to cortisone in the hair shaft (Yang et al., 1998). Cortisone is less polar than cortisol, and thus could potentially be more readily incorporated into the hair shaft. Additionally, systemic cortisol has been shown to be directly incorporated into hair as both cortisol and cortisone (Kapoor et al., 2018). Because of this, cortisone was included as an analyte in order to provide a more comprehensive representation of total glucocorticoid concentrations and provide a more accurate measurement of wellbeing. Dehydroepiandrosterone (DHEA) was also included as an analyte because, like cortisol, it is another main adrenal steroid that is released in response to activation of the HPA axis and is thought to modulate various effects of glucocorticoids such as immune function and neurogenesis (Buoso et al., 2011; Karishma & Herbert, 2002; Kamin & Kertes, 2017; Mehta et al., 2014). Due to the antiglucocorticoid actions of DHEA, it has been hypothesized that the cortisol:DHEA ratio may provide a more accurate reflection of overall HPA axis regularity with a lower ratio reflecting a healthier stress response (for review see Kamin & Kertes, 2017; Markopoulou et al., 2009; Mehta et al., 2014; Schury et al., 2017). We therefore included this ratio as an outcome measure of wellbeing for this study.

Together, the results of previous studies demonstrate that acute experience with ECE has short-term beneficial effects for NHP wellbeing and is associated with changes in salivary cortisol. However, both the effect of long-term experience with ECE on NHP wellbeing and the persistence of the effects from the use of ECE remain unknown. Furthermore, it is unclear how different social environments or life history may modulate the effects of ECE in NHP. The study reported here provides a preliminary examination of the effects of long-term ECE exposure on HPA hormones measured from the hair of adult rhesus macaques. In addition, the long-term effects of infant social environments were examined. The monkeys that participated in this study were either reared in a social group with their mothers (mother reared - MR) or in a social group consisting of only age-mate peers (nursery reared - NR) as infants before being united in a combined social group at approximately 7-months of age. The nursery rearing paradigm is often considered a model of early life adversity because of the disruption in the mother-infant attachment relationship (Conti et al., 2012). The effects of nursery rearing often parallel human affective disorders, with NR monkeys differing from their MR counterparts in many aspects, including behavior, physiology, and neurobiology (Bard & Hopkins, 2018; Capitanio et al., 2005; Chamove et al., 1973; Champoux et al., 1989; Conti et al., 2012; Dettmer et al., 2017; Harlow & Harlow, 1965; Machado & Bachevalier, 2003; Nelson & Winslow, 2009; Simpson et al., 2019; Spinelli et al., 2009; for review see Bennett & Pierre, 2010; French & Carp, 2016; Meyer & Hamel, 2014). Across age groups, NR monkeys exhibit behavior indicative of higher anxiety or reactivity compared to their MR counterparts (Champoux et al., 1991; see also Corcoran et al., 2012; see Dettmer & Suomi, 2014 for review). On the basis of these findings, the study reported here also examined whether differential rearing exerts long-term effects on hormonal measures associated with stress and whether experience with various types of enrichment modulates any effects.

Studies on the effect of differential rearing on the hypothalamic-pituitary-adrenal (HPA) axis have yielded mixed results (Table 1). Basal cortisol levels have been reported as higher in NR monkeys (Barrett et al., 2009), lower in NR monkeys (Shannon et al., 1998), or unaffected by rearing differences (Dettmer et al., 2009). Studies addressing the cortisol response to stressors in differentially reared macaques have yielded somewhat more consistent results, although discrepancies still exist. NR monkeys typically show a larger cortisol response to stressors compared to their MR counterparts (Dettmer et al., 2012; Fahlke et al., 2000; Feng et al., 2011; Higley et al., 1991; Higley et al., 1992). Conversely, two studies have found the opposite results, with MR monkeys showing a larger cortisol response to stressors (Capitanio et al., 2005; Clarke, 1993). However, to our knowledge nearly all studies examining cortisol in differentially reared macaques have looked at infant or juvenile monkeys or represent cross-sectional studies with sampling performed at multiple developmental timepoints. This is one of the first studies to examine hormone levels in hair in adult NHP from differential infant social rearing experiences. Additionally, this is the first study to examine the two rearing groups following ECE. We aim to fill this gap in knowledge to understand how early rearing experience interacts with exposure to ECE to influence long-term wellbeing.

**Table 1.**
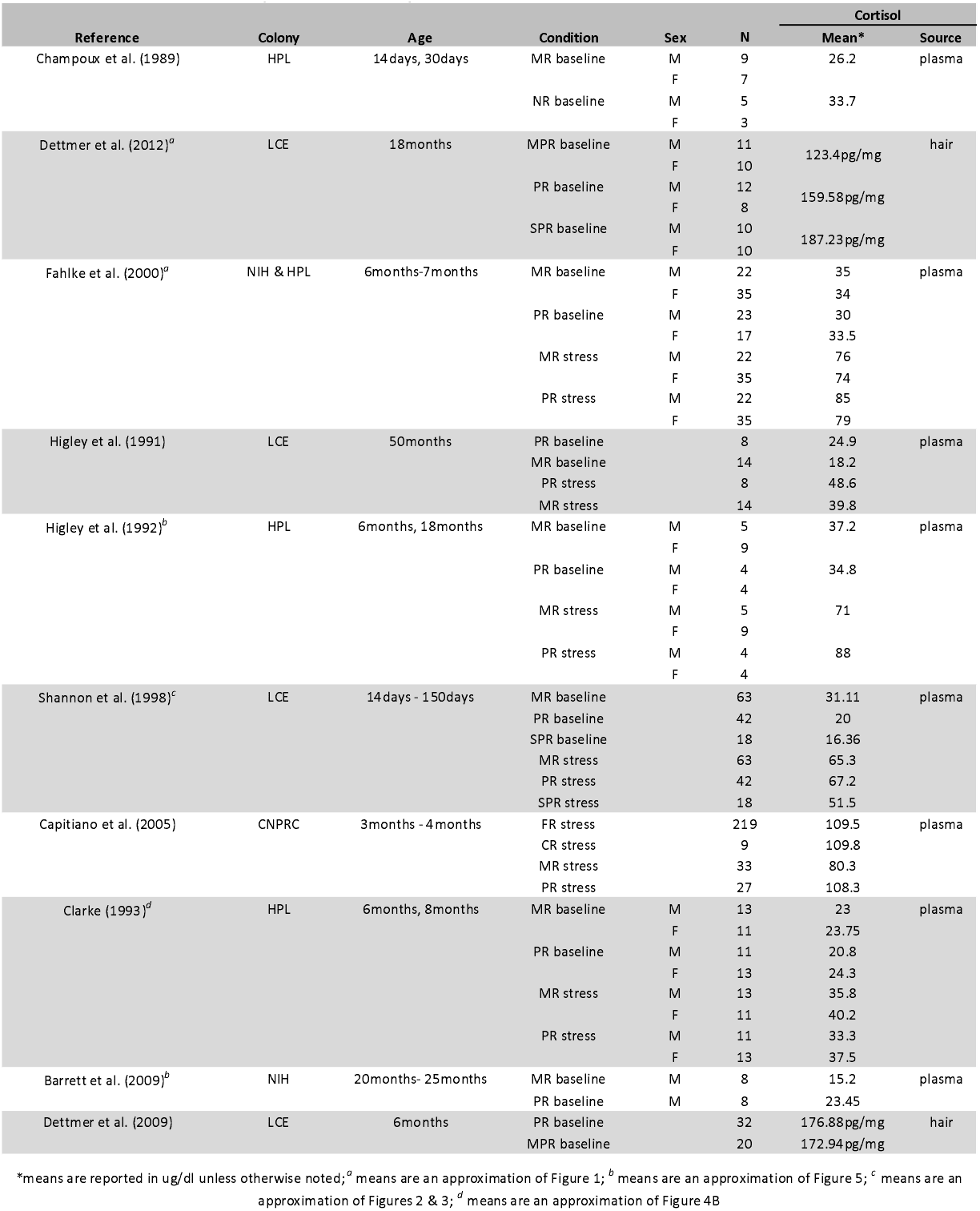
Cortisol Values from Differentially Reared Rhesus Macaques.

In the present study, we directly compared hair cortisol, cortisone, and DHEA from three groups of age-matched, adult male rhesus macaques with differential experience with ECE (CAU vs. ECE) and differential early social environments (MR vs. NR). The optimal fourth group (CAU-NR) was missing because of a lack of available subjects with appropriately comparable life histories that met these conditions. A secondary aim included examination of the long-term effects of differential rearing on hormone concentrations. We predicted that the ECE group would have lower cortisol, cortisone, and a lower cortisol:DHEA ratio than the CAU group that received care as usual enrichment and have had no experience with ECE. Further, we predicted that differential rearing could modulate this effect, with NR monkeys having higher cortisol and cortisone, and a higher DHEA:cortisol ratio than MR monkeys.

## 2. METHODS

### 2.1 Subjects

The sample was comprised of archival hair samples from 24 male rhesus macaques (*Macaca mulatta*) housed at either the Wisconsin National Primate Research Center (WNPRC; N=12) or the Harlow Primate Lab (HPL; N=12) at the University of Wisconsin-Madison. In total there were 12 animals in the care as usual group (CAU) who had never received videogame enrichment or other forms of enhanced cognitive enrichment (ECE), only standard environmental enrichment, and 12 animals who received regular ECE from one year of age through adulthood. This group had extensive experience with ECE through the use of the LRC-CTS both as a form of testing and enrichment. Ages at the time of hair samples ranged from 14.8 to 19.2 years of age (see Table 2). Within the ECE group, half of the adult monkeys had been reared with their mothers in infancy and half had been reared in peer-groups in a primate nursery (Nowak & Sackett, 2006; Shannon et al., 1998). Across both housing facilities subjects were maintained on a diet of monkey chow, fruits and vegetables, and *ad libitum* water as well as daily, care-as-usual enrichment. The standard enrichment generally included provision of fruits and vegetables, manipulanda (e.g., Kong toys, balls, nyla bones), foraging devices, perches, mirrors, and other common nonhuman primate environmental enrichment. All procedures, care, and housing were performed in accordance with care regulations of the participating facilities and their corresponding Animal Care and Use Committees.

**Table 2:** Subject characteristics.

| Group | N | Enrichment | Rearing | M age (years) ± SD |
| --- | --- | --- | --- | --- |
| CAU-MR | 12 | CAU | mother-reared | 16.3 ± 1.5 |
| ECE-MR | 6 | ECE | mother-reared | 18.7 ± 0.2 |
| ECE-NR | 6 | ECE | nursery-reared | 19.0 ± 0.2 |

This preliminary study relied on comparison of archival samples from colony animals and thus the animals’ histories were not tightly controlled. Consequently, animal records were evaluated in order to identify key aspects of the animals’ history with respect to influences on gonadal and adrenal hormones. One animal in the CAU-MR group was in a study on inhibition of gonadal and adrenal hormones three years prior to when hair samples were collected. Several of the animals had research history or were in active breeding at the time of sampling. In initial data analysis the hormone values for each animal were examined to determine whether they were outliers (+/- 3 standard deviations from the group mean; see below).

### 2.2 Materials. LRC-CTS

The apparatus that was used to present the videogame to the ECE group is described in detail in Bennett et al. (2016). Briefly, the device allows the joystick to be fixed to the monkeys’ enclosures while providing easy viewing of the screen and easy access to the reward pellet that is released when the monkey completes the activity. The monkey enclosures are slightly modified with a metal door (9 x 13cm), which can be removed to allow access to the joystick and reward. Monkeys in the ECE group have had experience with most LRC-CTS tasks, although most frequently the SIDES and CHASE tasks.

### 2.3 Hair collection

Hair samples from each group were obtained during routine semi-annual clinical health veterinary exams while the animals were anesthetized. Hair was taken from the intra-scapular region and cut at the proximal end near the follicle. Samples were kept in opaque aluminum foil pouches and stored at room temperature until assayed.

### 2.4 Assay method

Hair washing, steroid extraction, and LC/MS/MS followed the methods of Kapoor et al. (2014) with minor modifications. Hair was washed twice with 3mL of isopropanol and vortexed for 3min after each wash. After the second vortex, the hair was dried in polypropylene tubes in the dark until processing. Exactly 15mg of unground hair was then weighed out and placed into a clean culture tube. For liquid extraction, 2mL of methanol was added to each tube and tubes were incubated in the dark at room temperature for 48 hours. After incubation, internal standard, 1.5mL ammonium bicarbonate buffer pH 10, and 2mL 70:30 ethyl acetate hexanes were added to the tubes. Tubes were then vortexed for 8mins and centrifuged at 1500 rpm for 3min. The organic phase was pipetted off the top and transferred to a clean test tube. Samples were then dried down and reconstituted in 25ul 20% acetonitrile/water, vortexed well, and centrifuged at 1000 rpm for 1min. Reconstitute was then transferred into minivials and froze until analysis.

### 2.5 Data Analysis

Basic descriptive statistics were analyzed to check for normality. Because kurtosis values for some analytes were above two, all values were log transformed to assure normality and log base ten of analyte values were used for analysis. In order to examine the effect of experience with cognitive enrichment on hair analytes, a between-subjects MANOVA was run with analytes as the dependent variables and group as the independent variable. Analytes included cortisol + cortisone, DHEA, and cortisol:DHEA ratio. Cortisol and cortisone were summed for all statistical analyses for reasons described above. In order to test for equality of variance between groups, separate F-tests were run with analytes as the dependent variables and group as the independent variable. A regression with correlation coefficient was run with cortisol and DHEA in order to determine how cortisol and DHEA vary together in individuals between groups.

### 2.6 General Findings

Prior to analysis, examination of the data revealed that one animal in the CAU-MR group had values greater than 3 standard deviations higher than the mean for several analytes and thus was considered an outlier and excluded from analysis. No other animals were outliers and thus were all included in the analysis. An initial ANOVA indicated that age was significantly different between groups (*F*_2,20_ = 17.832, *p* < 0.0001). In order to control for this, age was kept as a covariate for all subsequent analyses.

## 3.0 RESULTS

### 3.1 CAU Enrichment vs ECE

Overall, the groups did not significantly differ as a function of enrichment condition (ECE vs CAU) in mean cortisol + cortisone, DHEA, or cortisol:DHEA ratio. Untransformed mean ± standard deviation analyte levels for each group are reported in Table 3. The equality of variance F-test revealed no significant difference in variability of log cortisol or log cortisol + cortisone between ECE-MR and CAU-MR groups. There was, however, a significant difference in variability of log DHEA between the ECE-NR and CAU-MR groups (*F_5,10_* = 0.092, *p* < 0.0092) (Figure 1B). We also found a significant difference in variability of log cortisol:DHEA ratio between the ECE-MR and CAU-MR groups (*F_5,10_* = 0.180, *p* < 0.0463), with this difference being even more significant between the ECE-NR and CAU-MR groups (*F_5,10_* = 0.147, *p* < 0.0286) (Figure 1C). Upon discovering that the equality of variance assumption was violated, we ran a Kruskal-Wallis non-parametric test. Results showed no significant difference in mean rank between groups in log cortisol (H(2) = 0.242, *p* = 0.8861), log DHEA (H(2) = 0.196, *p* = 0.9065), log cortisol + cortisone (H(2) = 0.274, *p* = 0.8178), or log cortisol:DHEA ratio (H(2) = 0.116, *p* = 0.9437). Regression with correlation coefficient revealed that DHEA was more closely correlated with cortisol in the ECE-MR group (*R^2^* = 0.606) than in the CAU-MR group (*R^2^* = 0.086) (Figure 2).

**Table 3:** Untransformed mean ± SD analyte values by group.

| Analyte (pg/mg) | CAU-MR | ECE-MR | ECE-NR |
| --- | --- | --- | --- |
| cortisol | 7.7 ± 3.6 | 6.9 ± 2.8 | 6.1 ± 1.4 |
| cortisone | 6.7 ± 2.8 | 7.5 ± 3.1 | 7.0 ± 2.2 |
| cortisol+cortisone | 14.3 ± 6.2 | 14.5 ± 5.8 | 13.1 ± 3.6 |
| DHEA | 4.4 ± 3.8 | 2.9 ± 1.5 | 2.3 ± 0.6 |
| cortisol:DHEA ratio | 2.8 ± 2.1 | 2.6 ± 0.8 | 2.7 ± 0.9 |

**Figure 1.**
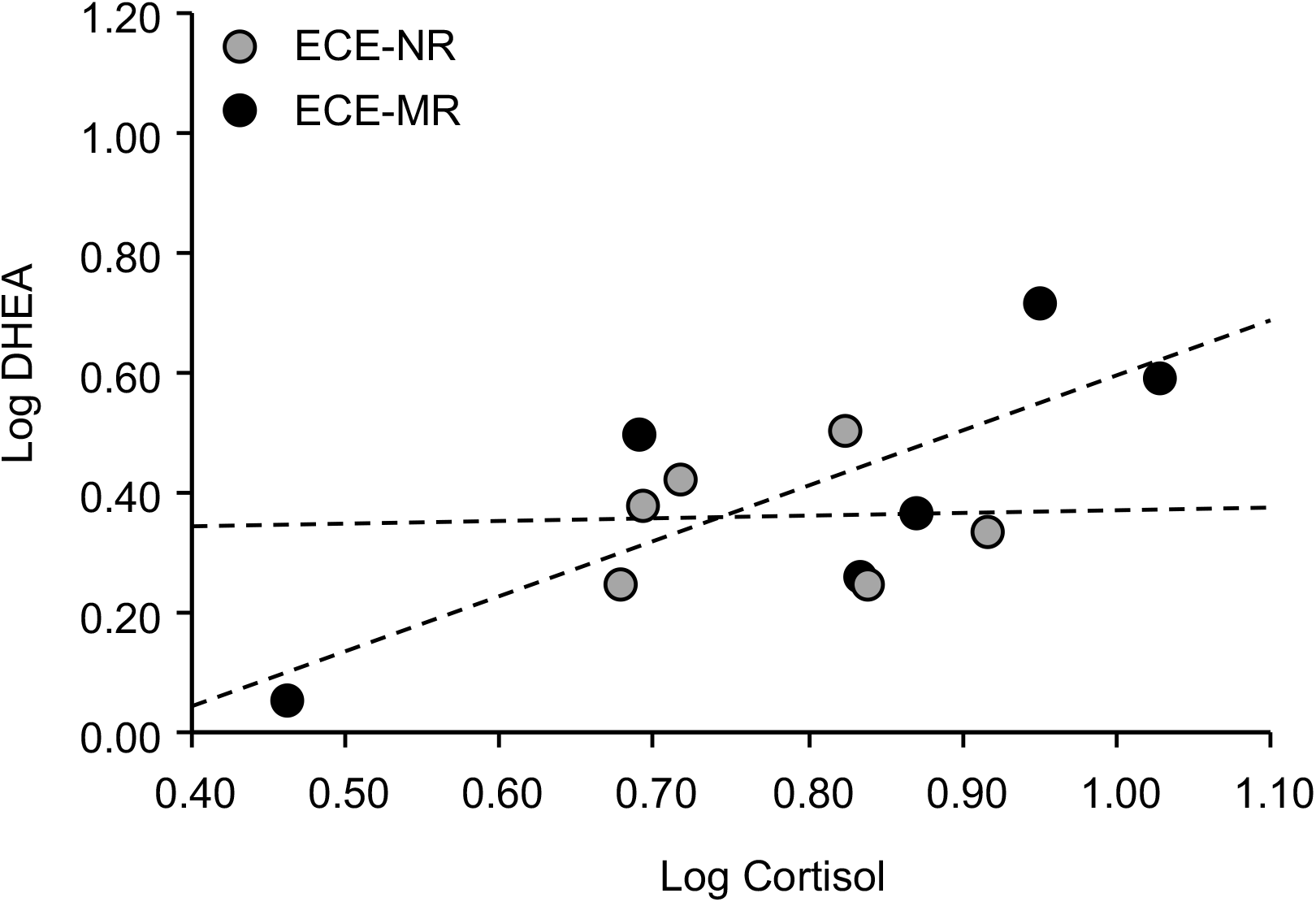
Comparison of variability of log analyte levels between groups. There were no significant differences in variability between the three groups in log cortisol+cortisone. The care-as-usual, mother-reared (CAU-MR) group showed significantly greater variability in log DHEA than the enhanced cognitive enrichment, nursery-reared (ECE-NR) group (*F_5,10_* = 0.092, *p* < 0.0092). The CAU-MR group also showed significantly greater variability in the log cortisol:DHEA ratio than both the ECE-NR group (*F*_*5*,10_ = 0.147, *p* < 0.0286) and enhanced cognitive enrichment, mother-reared (ECE-MR) group (*F_5,10_* = 0.180, *p* < 0.0463). *p<.05.

**Figure 2.**
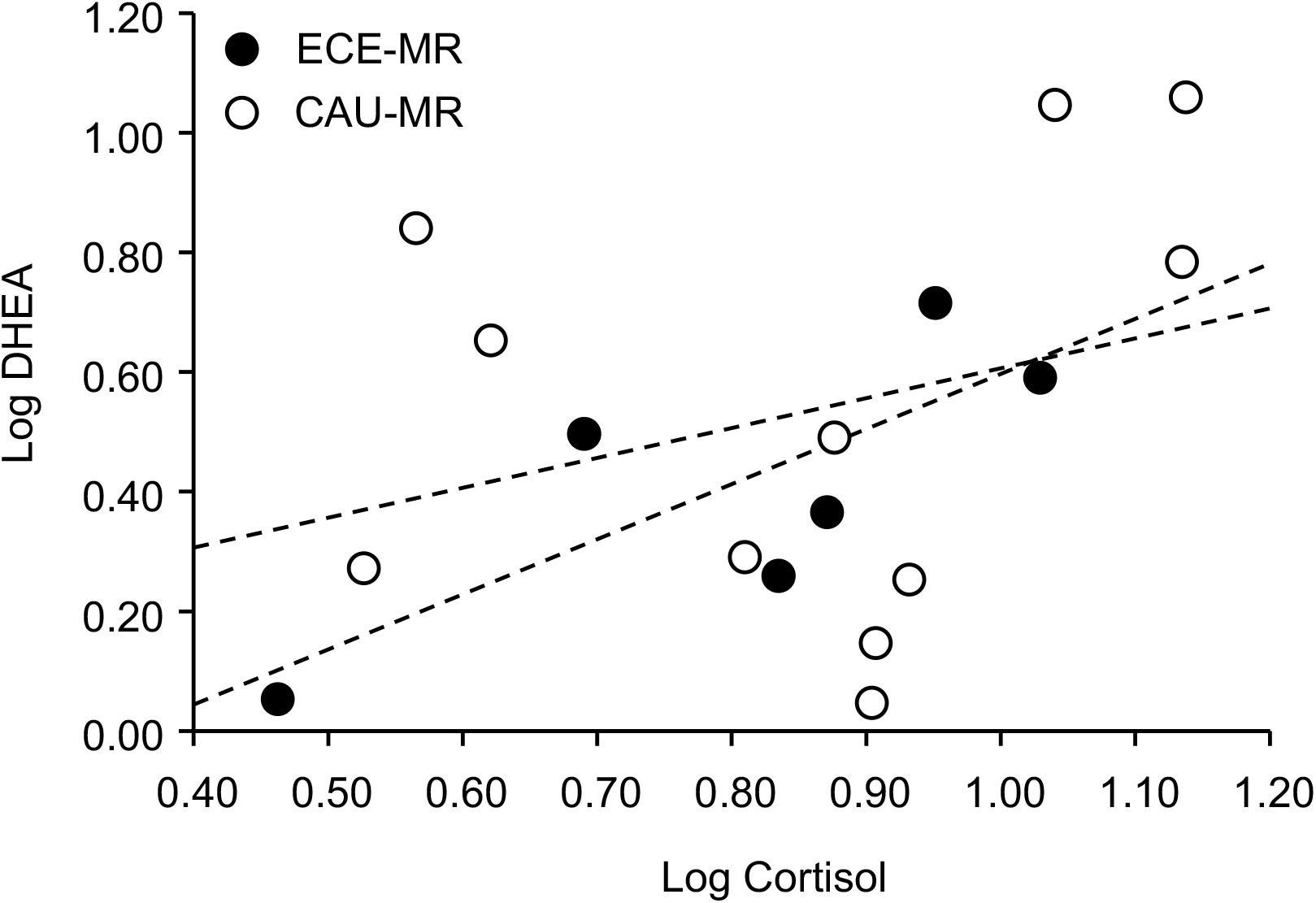
Log DHEA is plotted on the y-axis, while log cortisol is plotted on the x-axis in order to examine the relationship between the two analytes. Best fit lines are shown. Log DHEA and log cortisol are more closely correlated in the enhanced cognitive enrichment, mother-reared (ECE-MR) group *R^2^* = 0.606) than in the care-as-usual, mother-reared (CAU-MR) group (*R^2^* = 0.086).

### 3.2 Rearing

There were no significant differences in mean cortisol + cortisone, DHEA, or log cortisol:DHEA ratio between the ECE-MR and ECE-NR groups (Table 3). There were also no significant differences in variability of any of the analytes between these two groups, according to the equality of variance F-tests. Regression with correlation coefficient revealed that DHEA was more closely correlated with cortisol in the ECE-MR group (*R^2^* = 0.606) than in the ECE-NR group (Figure 3).

**Figure 3.**
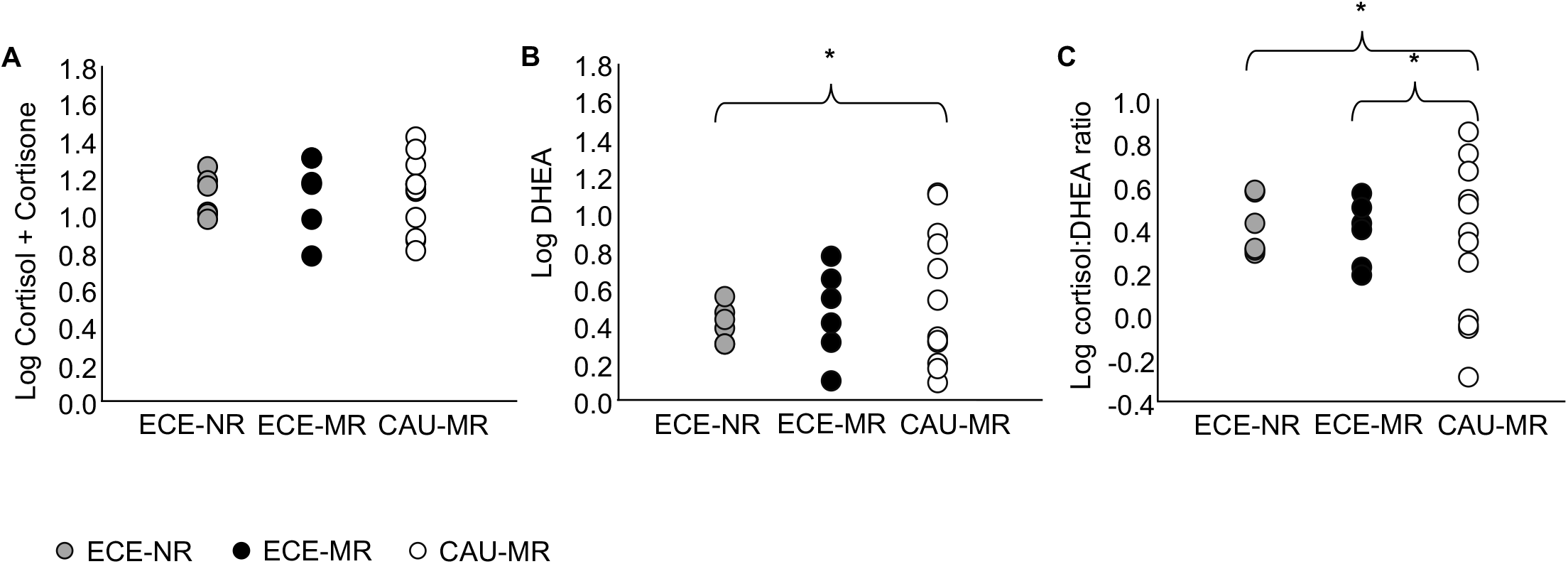
Log DHEA is plotted on the y-axis, while log cortisol is plotted on the x-axis in order to examine the relationship between the two analytes. Best fit lines are shown. Log DHEA and log cortisol are more closely related in the enhanced cognitive enrichment, mother-reared (ECE-MR) group (*R^2^* = 0.606) than in the enhanced cognitive enrichment, nursery-reared (ECE-NR) group (*R^2^* = 0.002).

## 4. DISCUSSION

Overall, the present preliminary study found no differences in mean cortisol + cortisone, cortisol:DHEA ratio, or DHEA between the three groups. There were, however, significant differences in variability between groups for the analytes DHEA and cortisol:DHEA ratio. The CAU-MR group had a higher degree of variance in cortisol:DHEA ratio than both the ECE-MR and the ECE-NR, which were not different from each other. The CAU-MR group had a higher degree of variance in DHEA than the ECE-NR group as well, however the CAU-MR group did not differ in variance from the ECE-MR group. Additionally, cortisol and DHEA varied together more strongly in both the ECE-MR and CAU-MR groups than in the ECE-NR group.

To our knowledge, this is the first study to provide preliminary analysis of hair cortisol levels following long-term experience with ECE. Our results suggest that experience with ECE has the potential to reduce inter-individual variability in stress hormones, as we observed in lower variation in DHEA and cortisol:DHEA ratio in the ECE groups compared to the CAU group. The finding that cortisol was more closely correlated with DHEA in the ECE-MR group compared to the CAU-MR group suggests that experience with ECE may also promote a healthier stress response than CAU practices alone. Overall, these results provide preliminary evidence consistent with a growing body of evidence that suggests ECE is beneficial to NHP wellbeing. Our observations are consistent with the findings of previous studies which demonstrated that acute experience with ECE lowers salivary cortisol and reduces undesired behaviors (Fagot et al., 2014). It remains for future study to confirm the result by directly examining whether experience with ECE influences reactivity to stressful situations.

To our knowledge, this is the first comparison of cortisol, cortisone, and DHEA levels in middle-aged rhesus macaques that were differentially reared in infancy. No significant differences were found in mean analyte values or variability between the ECE-MR and ECE-NR groups. This was unexpected because previous literature has reported differences in cortisol values between MR and NR monkeys in infancy and adolescence (see Table 1), however, we note that our sample size is small. In addition, we sampled adult rhesus macaques, rather than infants or adolescents and it is possible that rearing group differences are less apparent at older life periods. In contrast to the ECE-MR group, the correlation between cortisol and DHEA was almost nonexistent in the ECE-NR group. This pattern of results may suggest persistent effects of infant social experience, with ECE experiences serving as a form of intervention that modulated some of the adverse effects of early life stress on the HPA axis but was unable to correct the change in the relationship between DHEA and cortisol as it pertains to the stress response. Interpretation of this result is difficult because the relationship between DHEA and cortisol is not fully understood. Future studies could seek to clarify this hypothesis by comparing analyte values and correlations throughout the lifespan of differentially reared animals both with and without experience with ECE. Additionally, further research on the roles of DHEA, cortisol and their relationship in the stress response could further clarify our results.

The effectiveness of glucocorticoids as a measurement of wellbeing has been debated, and thus provides a limitation to the interpretation of our results. Although it is known that glucocorticoid synthesis increases in response to stressful situations, synthesis of these hormones can be influenced by a variety of other stimuli as well, some of which may not be decremental to welfare (Olsson & Dahlborn, 2002; McEwen, 1998). Additionally, the effects of glucocorticoids are influenced by many factors, such as the amount of free glucocorticoids and amount of glucocorticoid receptors (Ralph & Tilbrook, 2016). It is important to remember when interpreting our results that it is not the presence of glucocorticoids alone that influences animal welfare, but the actions of these hormones. Because numerous studies have previously demonstrated behavioral improvements in response to ECE, we are confident in our interpretation that provision of ECE improves NHP welfare (Bennett et al., 2016; Fagot et al., 2014). A secondary limitation to our study is the age of participants. It is known that hormone concentrations can decline with age, and as the monkeys that participated in this study were adults it is possible that this may have influenced hormone concentrations (Fourie & Bernstein, 2011). In order to control for any effects aging may have had on our study, participants were age-matched.

In conclusion, the observations described here suggest that although experience with ECE vs. CAU practices did not influence average cortisol, DHEA, or cortisol:DHEA ratio, monkeys that had experience with ECE had lower variability in hormone measurement, suggestive of a more typical stress response. In turn, our results add to a growing body of literature (Bennett et al., 2016; Mallavarapu et al., 2013; Perdue et al., 2017; Washburn & Rumbaugh, 1992a) that provide evidence for ECE as a form of intervention to promote healthy development in NHP and support its use as an effective form of enrichment in animals’ care.

## ACKNOWLEDGEMENTS

The authors gratefully acknowledge the assistance of Christopher Corcoran in the collection from rhesus monkeys and to the facility managers and animal care technicians at the Harlow Center for Biological Psychology. Observations with the animals were made in accordance with the Guide for the Care and Use of Laboratory Animals published by the US National Institutes of Health (NIH Publication No. 85-23, revised 2011). The research and manuscript preparation were partially supported by NIH grants MH084980 and P51OD011106.

